# Reducing individual differences in task fMRI with OGRE (One-step General Registration and Extraction) preprocessing

**DOI:** 10.1101/2023.09.19.558290

**Authors:** Mark P. McAvoy, Lei Liu, Ruiwen Zhou, Benjamin A. Philip

## Abstract

Volumetric analysis methods continue to enjoy great popularity in the analysis of task-related functional MRI (fMRI) data. Among these methods, the software package FSL (FMRIB, Oxford, UK) is omnipresent throughout the field. However, it remains unknown what advantages might be gained by integrating FSL with alternative preprocessing tools. Here we developed the One-step General Registration and Extraction (OGRE) pipeline to apply FreeSurfer brain extraction for simultaneous registration and motion correction (“one-step resampling”), for FSL volumetric analysis of fMRI data. We compared three preprocessing approaches (OGRE, FSL, and fMRIPrep) in a dataset wherein adult human volunteers (N=39) performed a precision drawing task during fMRI scanning. The three approaches produced grossly similar results, but OGRE’s preprocessing led to lower inter-individual variability across the brain and greater detected activation in primary motor cortex contralateral to movement. This demonstrates that FreeSurfer tools and one-step resampling can improve FSL’s volumetric analysis of fMRI data. The OGRE pipeline provides an off-the-shelf method to apply FreeSurfer-based brain extraction and one-step resampling of motion correction and registration for FSL analysis of task fMRI data.

## INTRODUCTION

Numerous methods and software packages exist for the analysis of fMRI data, and many of these approaches offer different advantages and tradeoffs. One commonly used analysis package is the fMRIB Software Library (FSL; Jenkinson et al., 2012), particularly its FEAT software for general linear model (GLM) volumetric model-based analysis of fMRI data (Woolrich et al., 2004; Woolrich et al., 2001). FSL is highly popular: as of June 2024, the reference for FSL’s Brain Extraction Tool (BET; Smith, 2002) has over 12000 citations, including over 1000 in the preceding eighteen months (Google Scholar). However, despite FSL’s popularity, FSL preprocessing has some shortcomings, one of which is its reliance on BET for identifying brain voxels in the scan volume. BET is prone to under- and over-extraction of brain voxels (Boesen et al., 2004; Klein et al., 2010; Zhuang et al., 2006), and has limited options for adjusting its algorithmic criteria (Jenkinson, 2018). Together these shortcomings mean that BET either produces suboptimal brain extractions or requires time-consuming and error-prone manual adjustment (Mohapatra et al., 2023; Quilis-Sancho et al., 2020). Therefore, alternative FSL-compatible preprocessing approaches (including brain extraction) could improve the reliability and accuracy of fMRI data analysis.

One alternative preprocessing approach takes advantage of the FreeSurfer parcellation (e.g. Fischl & Dale, 2000; Fischl et al., 2002; Fischl et al., 2004) to refine the brain extraction and improve the alignment, registration, and motion correction of functional data by means of a novel “one-step resampling” procedure with FSL tools (Glasser et al., 2013). This process removes spatial artifacts and distortions, creates cortical surface and myelin maps, performs precise within-individual cross-modal registrations, and carries out volume-based registrations to standard spaces. The one-step resampling procedure is a small portion of the full Human Connectome Project (HCP) processing pipeline, which subsequently uses a surface-based approach with numerous further advantages (Coalson et al., 2018; Dickie et al., 2019; Glasser et al., 2018; Glasser et al., 2016). However, volumetric approaches continue to enjoy widespread popularity as lower resolution fMRI data remains common in the field (e.g. Farahibozorg et al., 2021; Holmes et al., 2015), and volumetric approaches require less computing infrastructure than surface-based analysis. Few studies have assessed the potential advantages gained by incorporating the one-step resampling procedure into FSL volumetric analysis, because direct, controlled tests (i.e. controlling for later processing steps) have not been possible because the existing standard in FreeSurfer-based preprocessing, fMRIprep (Esteban et al., 2019), neither provides off-the-shelf appliation to FSL FEAT (Williams, 2019) nor includes the one-step resampling procedure.

Here, we created the OGRE (One-step General Registration and Extraction) pipeline to integrate the “one-step resampling” procedure as a general-purpose fMRI preprocessing tool for FSL volumetric analysis. We performed a fully-controlled comparison of three preprocessing pipelines: OGRE vs. fMRIprep vs. FSL preprocessing, by evaluating the effects of the three preprocessing pipelines on a fMRI motor task with the same subsequent volumetric FSL FEAT GLM analysis. The three preprocessing methods were compared for differences in task-based activation, brain extraction, and inter-individual variability.

## MATERIALS & METHODS

### Participants

Thirty-seven right-handed adults (28 female; ages 48 ± 18, range 24-82) performed a precision drawing task (see *fMRI Task* below) in the fMRI scanner. 12/37 participants had peripheral nerve injuries to their right arm, but the differences between groups (categorical or performance-related) were not a focus of the current study. All participants gave informed consent, and all procedures were approved by the Institutional Review Board at Washington University in St. Louis School of Medicine.

### fMRI Acquisitions

Scans were performed on a Siemens PRISMA 3T MRI scanner. BOLD EPIs for fMRI were collected using a T2*-weighted gradient echo sequence, a standard 64-channel birdcage radio-frequency coil, and the following parameters: TR = 662 ms, TE = 30 ms, flip angle = 52°, 72 x 72 voxel matrix, FOV = 216 mm, 60 contiguous axial slices acquired in interleaved order, resolution: 3.0 x 3.0 x 3.0 mm, bandwidth = 2670 Hz/pixel, multi-band acceleration = 6x. Siemens auto-align was run at the start of each session.

High-resolution T1-weighted structural images were also acquired, using the 3D MP-RAGE pulse sequence: TR = 4500 ms, TE = 3.16 ms, TI = 1000 ms, flip angle = 8.0°, 256 x 256 voxel matrix, FOV = 256 mm, 176 contiguous axial slices, resolution: 1.0 x 1.0 x 1.0 mm. A T2-weighted image was also acquired at: TR = 3000 ms, TE = 409 ms, 256 x 256 voxel matrix, FOV = 256 mm, 176 contiguous axial slices, resolution: 1.0 x 1.0 x 1.0 mm. Spin echo field maps were also collected before the functional runs.

### fMRI Task

Each participant completed 3 BOLD functional scans. During each scan, the participant used their right hand to perform a precision drawing task based on the STEGA app (Philip & Frey, 2014; Philip et al., 2023). (These runs were interleaved with 3 additional runs where the participants used their left hand to perform the same task, but these left hand runs are not included in the current study.) also used their left hand Participants saw hollow shapes, and were instructed to draw a line within the bounds of the shape, as fast as possible while prioritizing staying in-bounds over speed. Participants saw a video overlay of their (transparent) hand/pen over the drawn shape, using an MRI-compatible drawing tablet with bluescreen technology (Karimpoor et al., 2015), as shown in **Figure 1**. The task was presented in a block design with 15.2 seconds (23 images) of drawing, followed by 15.2 seconds (23 images) of rest (fixation cross). A scan comprised ten cycles of draw/rest, with an additional rest block at the start, and 3.3 seconds (5 images) of additional rest after the final rest block, leading to a total duration of 5:23 (230 images of drawing, 258 rest) per scan.

**Figure 1:**
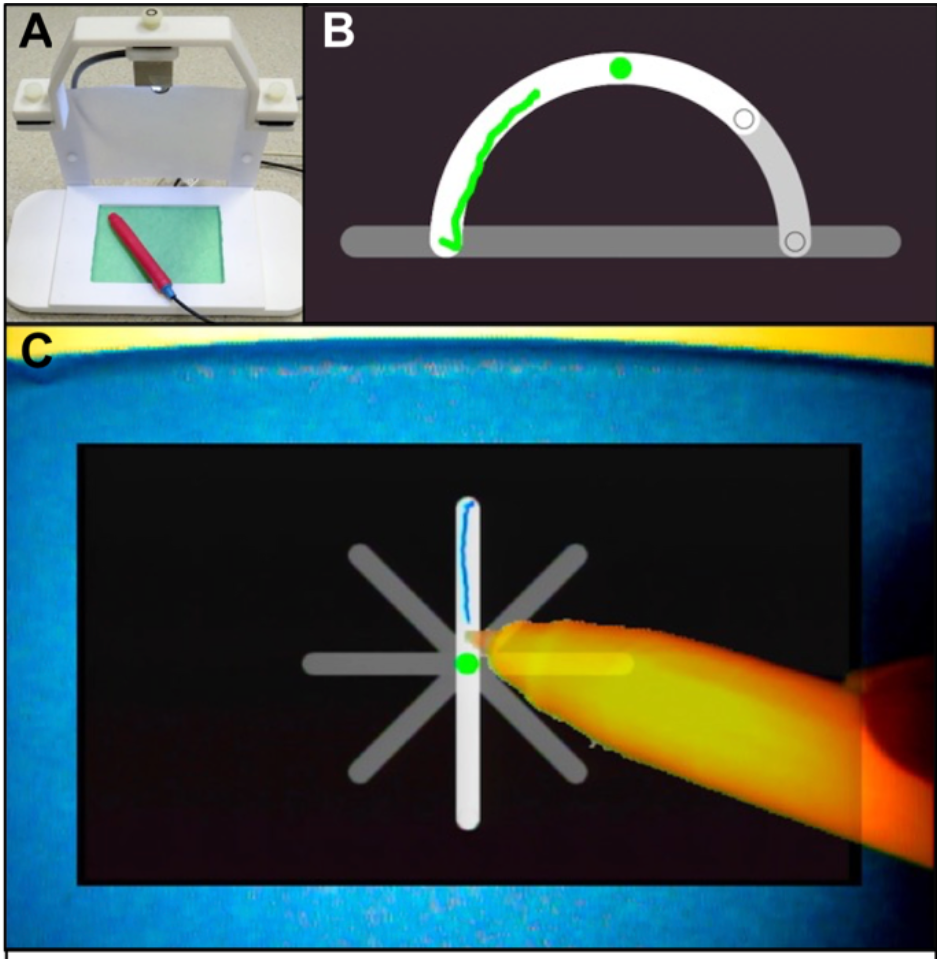
Motor task used for pipeline testing. A: MRI-compatible tablet. B: STEGA precision drawing task. C: Participant view of bluescreen integration.

### Data Preprocessing

#### One-step General Registration and Extraction (OGRE) Pipeline

OGRE (One-step General Registration and Extraction) was developed on Macintosh (Apple, Cupertino CA) and is based on version 3.27 of the HCP pipeline (https://github.com/Washington-University/HCPpipelines/tree/v3.27.0).

Preprocessing steps include FreeSurfer parcellation and brain extraction, motion correction, field map distortion correction, and warping to the 2mm MNI atlas via the “one-step resampling” via FSL FLIRT and FNIRT (Andersson et al., 2007) as detailed in Glasser et al. (2013). New features added for OGRE included the option to specify native resolution as 0.7, 0.8, or 1.0 mm; the option to leave fMRI time series in native HCP space instead of registering to MNI atlas; extended compatibility to FreeSurfer versions 7.2.0, 7.3.2, 7.4.0 and 7.4.1 (https://surfer.nmr.mgh.harvard.edu/); adjustable dilation/erosion settings for brain masks; and the addition of three new steps to match FEAT: optional nonlinear spatial filtering (Smith & Brady, 1997), optional nonlinear temporal high pass filtering (Woolrich et al., 2001), and a switch from mean-based to median-based intensity normalization. OGRE also produces FEAT-formatted registration images and matrices for higher-level FEAT analysis. OGRE preprocessing outputs were entered as data for analysis in FEAT using its “Statistics” option (rather than “Full Analysis,” since OGRE has replaced FEAT preprocessing).

OGRE software and documentation are available online at: https://github.com/PhilipLab/OGRE-pipeline

### Parallel analyses

#### OGRE vs. fMRIprep vs. FSL-only

In the “OGRE analysis,” the OGRE pipeline described above was used to preprocess data with the following options: FreeSurfer 7.4.1, native resolution of 1 mm, distortion correction using spin echo field maps, spatial smoothing kernel of 6 mm FWHM, intensity normalization via “grand median scaling” to a value of 10000, high pass temporal filtering with 60 sec cutoff, and output registration to 2 mm MNI atlas. Subsequently, GLM statistical analysis was performed with FEAT version 6.0.7 (Smith et al., 2004). Explanatory variables (EVs) were modeled, along with their temporal derivatives, according to the block design described in the previous section (*fMRI Task*), with additional confound EVs based on head motion parameters (translation and rotation). Volumes with excess head motion were addressed (Siegel et al., 2014) via additional confound EVs calculated within each time series as framewise displacement of 75^th^ percentile + 1.5* interquartile range, using the FSL script FSLMotionOutliers. The hemodynamic response was accounted for by convolving the model with a double-gamma function. First-level contrasts of parameter estimates (COPEs) were calculated for Task vs. Rest. The first-level COPE (Task > Rest) served as input to higher-level analyses performed using a fixed-effects model. Z-statistic (Gaussianized T) images were thresholded using clusters determined by Z ≥ 3.1 and a corrected cluster significance threshold of p < 0.05. The first-level COPEs were averaged across runs for each participant (second level). Region of interest (ROI) analyses were performed on second-level (i.e. participant-level) data, as detailed in *ROI Analyses* below.

In the “fMRIprep analysis,” preprocessing was performed using fMRIprep 23.2.3 (Esteban et al., 2019), using default settings except that slice timing was ignored, sub-millimeter reconstruction option was disabled, and outputs were registered to 2mm MNI atlas. Full fMRIprep methods are described in **Supplementary Text**. FSL was used to perform spatial smoothing with a 6 mm FWHM and high pass temporal filtering with 60 sec cutoff, and registration outputs were modified following established workarounds for integrating fMRIprep output with FSL (Mumford, 2017). Subsequently, GLM statistical analysis was performed with FEAT as described for the OGRE analysis above.

In the “FSL-only analysis,” preprocessing included the following steps: Non-brain structures were removed using BET. Head movement was reduced using MCFLIRT motion correction. Distortion correction was applied using spin echo field maps using FSL PRELUDE and FUGUE. Paralleling the OGRE implementation, spatial smoothing, intensity normalization, and the high-pass temporal filter were applied with the same settings as above. Functional data were registered with the high-resolution structural image using boundary-based registration (Greve & Fischl, 2009), and resampled to 2x2x2 mm resolution using FLIRT; the participant images were then registered to standard images (Montreal Neurological Institute MNI-152) using FNIRT nonlinear registration (Andersson et al., 2007). Subsequently, GLM statistical analysis was performed with FEAT as described for the OGRE analysis above.

#### ROI Analyses

To quantitatively compare the three analyses, a region-of-interest (ROI) approach was used, using an atlas of 300 volumetric ROIs with known network assignments (Seitzman et al., 2020). This atlas assigned each area to one of 14 networks (auditory, cingulo-opercular, default mode, dorsal attention, frontoparietal, medial temporal, parietomedial, reward, salience, somatomotor dorsal, somatomotor lateral, ventral attention, and unassigned). For each participant, the signal magnitude was calculated as the mean % BOLD signal change for the Task > Rest contrast in second-level (within participant) analyses.

The three analyses were compared via two approaches. The first approach focused on inter-individual variability. The primary variable was the standard deviation (across the signal magnitudes of the 39 participants, within each ROI). Direct statistical comparisons within each ROI used a pairwise Pitman-Morgan test (Morgan, 1939) to compare the inter-individual variability of OGRE signal magnitudes vs. the inter-individual variability of FSL-only signal magnitudes. To explore the spatial patterns of where the two analyses differed, a 3 (Method: OGRE, fMRIprep, FSL-only) * 14 (Network) * 2 (Hemisphere: left, right) ANOVA was performed on the standard deviations of the network ROIs. One ROI (in visual cerebellum) was omitted from the ANOVA because it was not associated with either hemisphere, leaving 299 ROIs in the ANOVA. Post-hoc tests were performed using Tukey’s Honestly Significant Difference procedure for effects of interest (i.e. involving the Method factor) at alpha = 0.05.

The second approach for comparing analyses focused on mean signal magnitude. The same process was followed as above, except: the primary variable was the mean signal magnitude (across the 39 participants). Direct statistical comparisons used paired-sample t-tests.

Separately from the above, task-specific results were compared across analyses by quantifying results in a normatively defined left hemisphere hand motor cortex ROI, (Smith & Frey, 2011) a sphere of 5 mm radius centered on MNI coordinates X = -38, Y = -24, Z = 54. This ROI was used to compare the results from the three analyses by measuring each analysis’ Z-score for Draw > Rest at the top level (across participants). Z-score distributions were compared with paired-sample t-tests, α = 0.0167 (0.05 / 3 to provide Bonferroni correction for three pairwise comparisons).

#### Whole-brain analyses

To evaluate whole-brain results from each pipeline, a third-level analysis was performed to identify the mean activation (for Draw > Rest) within each pipeline. Subsquently, the three pipelines were compared via repeated-measures FEAT GLMs treating the analyses as multiple samples. Three pairwise comparisons were performed (i.e. one for each pair of preprocessing methods), with significant clusters were detected by Z ≥ 3.1 and a corrected cluster significance threshold of α = 0.0167.

#### Brain extraction and distortion correction comparisons

To quantify the difference in brain extraction and registration results between analysis methods, we compared the size of post-registration (i.e. standard space) brain masks from each task fMRI scan, following Zhang et al. (2019). Specifically, for all three preprocessing pipelines, comparisons were performed on the reg_standard/mask.nii.gz files generated for the FEAT GLM. Mask sizes were compared with other results with Pearson correlations, α = (0.05/3 to provide Bonferroni correction for 3 pairwise comparisons:

Differences between analyses in EPI distortion correction were not evaluated because all three analysis pipelines perform distortion correction via the same process. Specifically, in OGRE, EPI images are unwarped with the HCP implementation of the FSL’s topup algorithm (Andersson et al., 2007; Glasser et al., 2013; Smith et al., 2004). FSL’s topup is also currently used in fMRIprep when fieldmap images are available (Esteban et al., 2019), so fieldmap-based EPI unwarping should be identical across all three analysis pipelines. Fieldmap acquistions were available for all participants in this study.

## RESULTS

### Lower inter-individual variability with OGRE compared to FSL-only and fMRIprep analyses

OGRE analysis led to lower inter-individual variability (standard deviation) in most ROIs, with lower SD for OGRE compared to either FSL-only (z = -2.86, p = 3.1×10^-6^) or fMRIprep (z = 1.80, p = 3.2×10^-7^), as detailed in **Figure 2** and summarized in **Table 1**. For a full list of results for each of the 300 ROIs, see **Supplementary Table 1**.

**Table 1:** How often inter-individual variability differed between methods. FMP = fMRIprep. Counts are out of 300 ROIs. “Numeric advantage” defined as numerically lower standard deviation. “Sta0s0cal advantage” defined via Pitman-Morgan test, alpha = 0.05.

|  | Numeric advantage | Statistical advantage (method A) | Statistical advantage (method B) |
| --- | --- | --- | --- |
| OGRE vs FSL-only | OGRE 92.3% (277) | OGRE 80.3% (241) | FSL 2.0% (6) |
| OGRE vs fMRIprep | OGRE 89.0% (267) | OGRE 73.0% (219) | FMP 4.0% (12) |
| FMP vs FSL-only | FMP 56.3% (169) | FMP 11.7% (35) | FSL 19.0% (57) |

**Figure 2:**
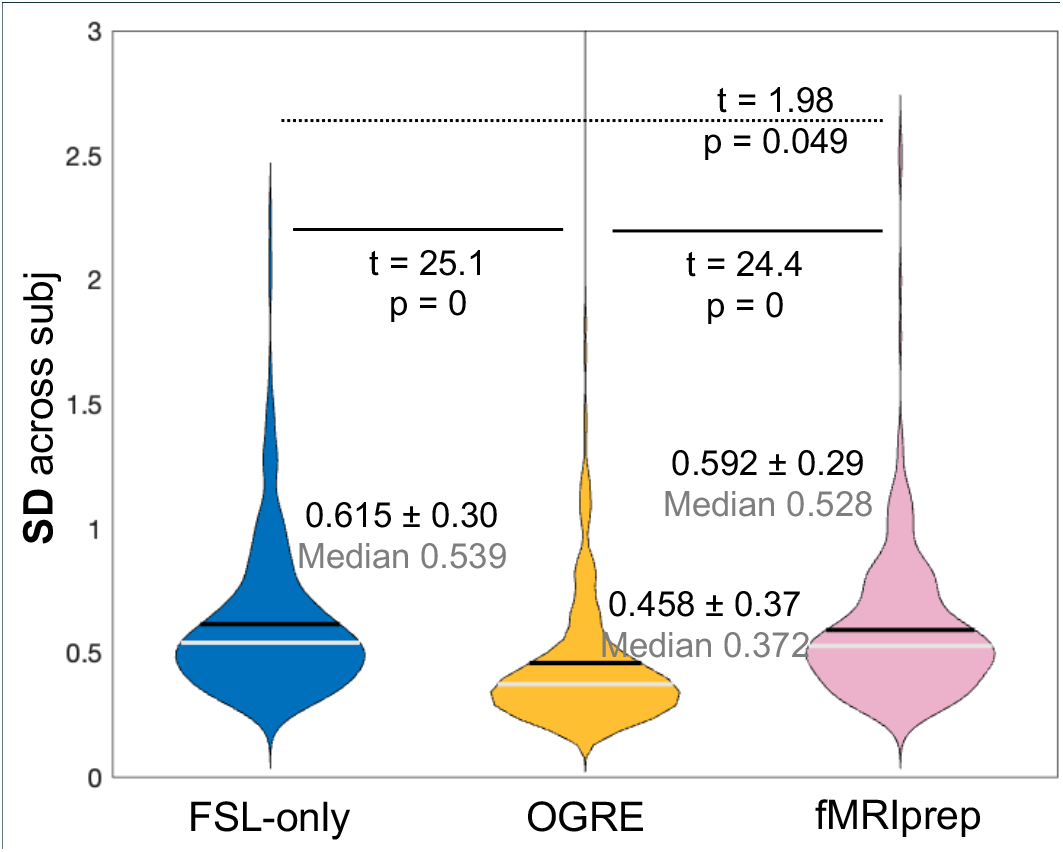
OGRE leads to lower inter-individual variability compared to FSL-only or fMRIprep. Each sample is a **SD** value (across 37 participants); each violin is the distribution of samples (across 300 ROIs). P-values from Pitman-Morgan test. Note that FSL-only vs. fMRIprep is not significant due to α = 0.0167.

To explore the pattern of where the three analyses differed, we performed a 4-way ANOVA: 3 (Method: OGRE, FSL-only, FSL-only) * 14 (network) * 2 (hemisphere). We found significant effects of method (F(2,836) = 5.9, p = 0.003) and network (F(13,836)=11.6, p = 2.3×10^-23^) but not hemisphere (p = 1.0). We found no significant interaction effects (p > 0.18). Post-hoc tests revealed that the network effect arose from higher variability for the networks Unassigned and Visual than most (7-10/14) networks. Critically, post-hoc tests revealed that the Method effect arose from lower inter-individual variability for OGRE (0.458 ± 0.370, median 0.372) than fMRIprep (0.592 ± 0.293, median 0.528) or FSL (0.615 ± 0.300, median 0.539).

### Minimal differences in BOLD signal magnitude between OGRE and other analyses

OGRE analysis led to higher signal magnitude (participant mean % BOLD signal change for Task > Rest) more often than lower signal magnitude, compared to FSL-only analysis (t = -3.9, p = 1.2×10^-4^); and fMRIprep led to higher signal magnitudes than FSL-only analysis (t = -9.75, p = 1.1×10^-19^), as detailed in **Figure 2** and summarized in **Table 2**. For a full list of results for each of the 300 ROIs, see **Supplementary Table 2**.

**Table 2:** How often mean signal magnitude differed between methods. FMP = fMRIprep. Counts are out of 300 ROIs. “Numeric advantage” defined as numerically lower magnitude. “Statistical advantage” defined via Wilcoxon Signed rank (nonparametic paired samples), alpha = 0.05.

|  | Numeric advantage | Statistical advantage (method A) | Statistical advantage (method B) |
| --- | --- | --- | --- |
| OGRE vs FSL-only | OGRE 61.3% (184) | OGRE 22.3% (67) | FSL 21.0% (63) |
| OGRE vs fMRIprep | OGRE 52.0% (156) | OGRE 22.0% (66) | FMP 23.0% (69) |
| FMP vs FSL-only | FMP 74.0% (222) | FMP 22.3% (67) | FSL 45.3% (136) |

To explore the pattern of where the three analyses differed, we performed a 4-way ANOVA: 3 (Method: OGRE, FSL-only, FSL-only) * 14 (network) * 2 (hemisphere)) on mean signal magnitudes. We found significant effects of Network (F(13,836) = 3.0, p = 1.7×10^-66^), but not Hemisphere (p = 1.0) or Method (p = 0.838). We found a significiant interaction effect for Hemisphere * Network (F(13,848) = 3.54, p = 2.1×10^-5^) but no others (p > 0.5). Post-hoc analyses revealed that the Network effect arose from higher mean activity in the Dorsal Attention network than all other networks, and higher in CinguloOpercular and SomatoMotor-Dorsal than in most (9/14) networks; and that the Hemisphere * Network interaction arose because the Salience network had higher mean activity in the left hemisphere than right hemisphere. However, we found no effects of Method on BOLD signal magnitude.

### Increased signal in in task-specific areas when using OGRE, especially vs. FSL-only

To assess the effects of OGRE analysis on the whole brain, we conducted whole-brain repeated measures analyses, treating the three analyses (OGRE, fMRIprep, FSL) as separate measures for each participant. Each analysis method led to grossly similar results, but with a scattered network of areas that showed significant differences between methods, as shown in **Figure 4A**.

To provide a quantitative contrast between analyses in an area with known “ground truth,” we compared results across methods in left hemisphere normative hand M1, contralateral to the drawing right hand (**Figure 4B**). Z-scores for OGRE analysis were significantly higher than FSL-only (t = -3.51, p = 0.001), but non-significantly higher than fMRIprep (t = 1.74, p = 0.90); the difference between FSL and fMRIprep was not significant after multiple comparison correction α = 0.167 (t = -2.08, p = 0.45). While all three methods were perfectly capable of detecting task-specific activity in a task-relevant brain area, OGRE analysis was significantly better than FSL-only at detecting this task-specific activity, and produced the highest z-scores overall.

### OGRE identified smaller brain volumes, primarily through removal of non-brain voxels

To quantify the difference in brain extraction results between analysis methods, we compared the size of post-registration (i.e. standard space) brain masks of task fMRI runs. OGRE masks were substantially smaller than masks produced by other methods, incorporating fewer non-brain voxels, as detailed in **Table 3**. However, the importance of mask size depends on the method, as shown in **Table 4**. In FSL-only and fMRIprep analyses, results did not correlate significantly with mask size, suggesting that their large mask sizes did not impede analysis (likely due to the inclusion of distal non-brain areas with no BOLD signal). However OGRE performance was positively correlated with: Z-scores in contralateral M1 were significantly correlated with the number of total voxels and non-brain voxels, suggesting that OGRE produced masks that were too small (i.e. removed too many non-brain voxels for optimal performance under spatial smoothing), and OGRE – despite producing the highest Z-scores of the three methods – might achieve still better performance with larger brain estimates.

**Table 3:** Sizes of brain masks after registration to MNI-152 standard space.

|  | Total Voxels |  |  | Non-Brain Voxels |  |  |
| --- | --- | --- | --- | --- | --- | --- |
| Method | Mean ± SD | Median | Range | Mean ± std | Median | Range |
| FSL-only | 311,626 ± 8,952 | 312,174 | 287,183-332,575 | 83,741 ± 8,705 | 84,908 | 62,397-104,104 |
| OGRE | 266,321 ± 3,605 | 266,784 | 258,335-274,525 | 39,513 ± 3,227 | 39,671 | 32,637-47,245 |
| fMRIPrep | 387,073 ± 31,841 | 384,818 | 285,433-467,113 | 160,182 ± 32,209 | 157,555 | 63,280-239,357 |

**Table 4:** Relationship between brain mask size and analysis outcomes.

| Correlation between... | # voxels and BOLD signal |  | # voxels and M1 z-score |  | # non-brain voxels and BOLD signal |  | # non-brain voxels and M1 z-score |  |
| --- | --- | --- | --- | --- | --- | --- | --- | --- |
| Method | r | p | r | p | r | p | r | p |
| FSL-only | 0.043 | 0.799 | -0.063 | 0.709 | 0.034 | 0.84 | -0.083 | 0.623 |
| OGRE | 0.173 | 0.307 | 0.400 | **0.014** | 0.161 | 0.342 | 0.383 | **0.019** |
| fMRIPrep | -0.042 | 0.806 | 0.117 | 0.491 | -0.063 | 0.712 | 0.088 | 0.604 |

**Table 4:**
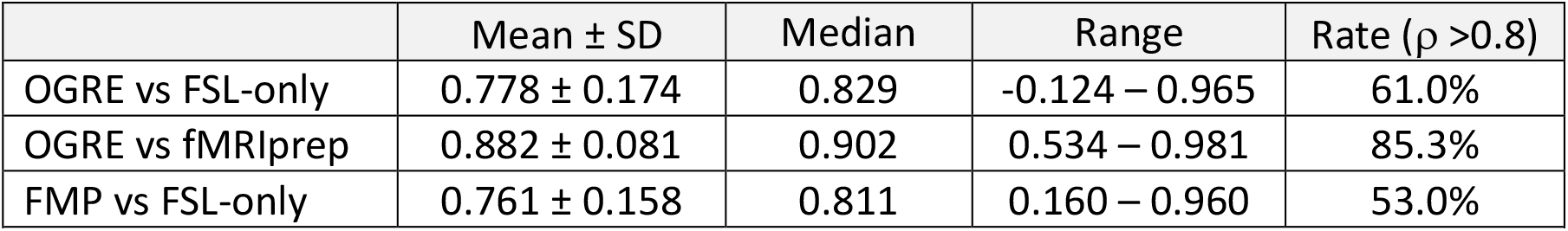
Correlations between methods, across 300 ROIs. Values = Spearman ρ.

### Differences between analysis methods are broadly consistent across participants

Our participant group included at least one major unmodeled source of variance: the distinction between typical adults and patients with peripheral nerve injury. This provides an advantage for practical testing, because it highlights OGRE’s robustness in the face of complex real-world data. However, we must rule out the possibility that this sample choice distorted our results. To address this, we first assessed whether participant group (patient vs. control) affected our measure of task-related BOLD signal (% BOLD signal change during Task > Rest). We performed a between-groups t-test to assess group differences in BOLD signal magnitude, and repeated this analysis separately for each ROI*analysis method (i.e. 900 comparisons), without multiple comparison correction. We found group effects in 13/300 ROIs with FSL analysis, 12/300 ROIs with OGRE, and 8/300 ROIs with fMRIprep, rates which do not significantly differ across analysis methods (for the most-different pair of methods, 𝒳^2^ = 1.2, p = 0.267). Therefore, peripheral nerve injury seemed to have no systematic effect on the measures tested here.

To further address the underlying concern about between-methods effects potentially being unevenly distributed across individuals, we examined the correlation of signal intensities (% BOLD signal change, Task > Rest) between methods, for each ROI. Correlations were high, especially for OGRE vs. fMRIprep, as shown in **Table 4**, showing a broadly linear relationship across methods, which suggests that the effects here do not depend on subgroups or systematic differences between participants.

## DISCUSSION

OGRE preprocessing led to greater detection of task-related activity and reduced inter-individual BOLD variability in volumetric FSL analysis of a motor task, which demonstrates the advantages of combining motion correction and registration via the “one-step resampling” process. OGRE had weaker advantages over fMRIprep analysis than FSL-only advantages, demonstrating separate benefits of Freesurfer brain extraction (shared by fMRIprep and OGRE) and motion correction via one-step resampling (OGRE only). The three methods produced grossly similar results, but for metrics with a clear definition of “better,” OGRE results were always significantly or trending better than all other methods. OGRE software will allow any FSL user to take advantage of these preprocessing benefits.

### Increased task-related signal in functional analysis after one-step resampling

We directly compared OGRE preprocessing with FSL and fMRIprep preprocessing by analyzing the same data with the same subsequent volumetric GLM analysis (in FSL FEAT). Our ultimate finding was that OGRE led to greater detection of BOLD signal in a brain area where activity is certainly task-relevant (M1 contralateral to movement, **Figure 4**). This detection of task-relevant activity (i.e. M1 Z-score) was significantly higher for OGRE than FSL-only; detection did not differ significantly between OGRE and fMRIprep, but OGRE had higher mean and median Z-scores than fMRIprep, indicating that OGRE’s signal detection performance was as-good-or-better versus any other method.

OGRE also led to increased BOLD signal in deep structures that likely represent noise (e.g. white matter, ventricles; **Figure 4A**). This may reflect amplification of real artifacts present in the data (e.g. movement-related). While amplification of artifacts is unhelpful, it is easily distinguishable from the expected cortical areas and networks involved in a task (here, precision drawing), and thus the heightened signal in deep structures should not present an obstacle to interpretation of task fMRI results.

Because we used real data rather than simulated data, it is impossible to determine whether increased signal in task-relevant areas reflects a more-precise measurement of a strong signal, or a less-precise measurement of a weak signal. However, the latter interpretation – artifactual inflation of BOLD signal strength – is incompatible with our finding that OGRE did not produce the highest signal magnitudes of our methods (**Figure 3**), but did produce the lowest inter-individual variability, as discussed in the next section.

**Figure 3:**
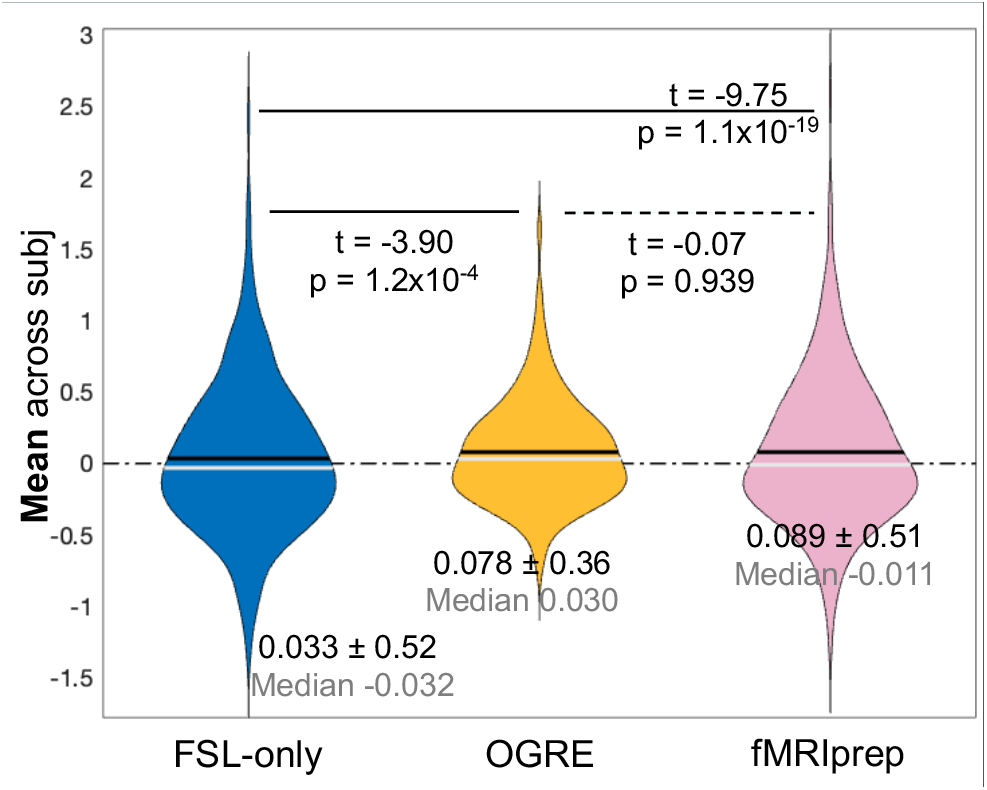
OGRE and fMRIprep lead to marginally higher mean activity than FSL-only. Each sample is a **mean** value (across 37 participants); each violin is the distribution of samples (across 300 ROIs). P-values from paired-samples t-test.

**Figure 4:**
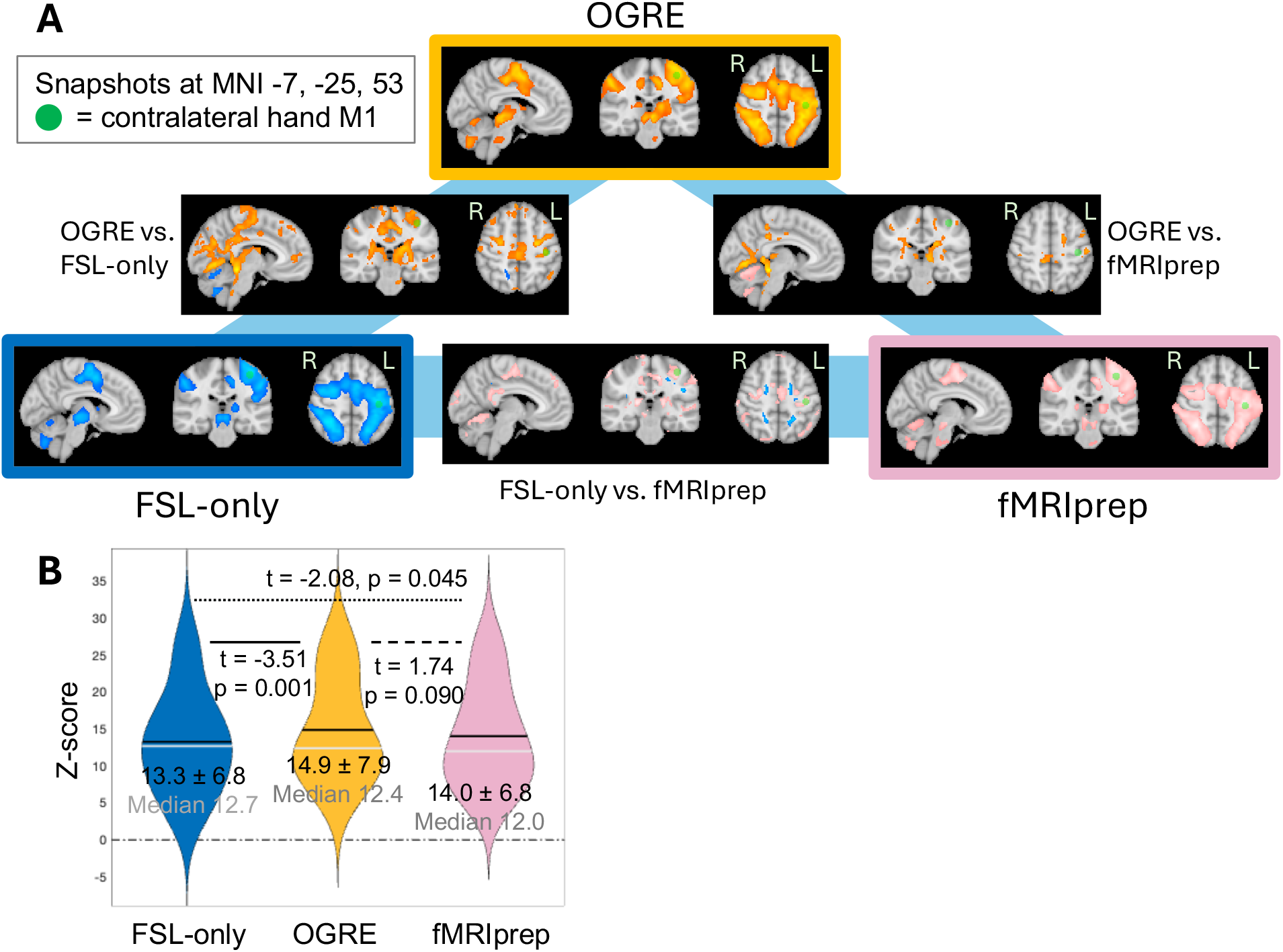
OGRE, FSL-only, and fMRIprep analyses lead to grossly similar activation maps during right hand drawing task, but OGRE produces strongest statistical results. **A:** Snapshot of whole-brain third-level (across participants) results for each analysis and between-analyses contrast. **B:** Quantitative comparison of statistical results in contralateral hand M1 (green circle in A). P-values from paired-samples t-test. Note that FSL-only vs. fMRIprep is not significant due to α = 0.0167.

### Reduced inter-individual variability after one-step resampling

OGRE preprocessing led to signigicantly lower inter-individual variability in BOLD signal across the brain, compared to either FSL or fMRIprep preprocessing (**Figure 2**). This is likely due to the two primary differences between OGRE and fMRIprep: OGRE’s one-step resampling procedure, or OGRE’s FNIRT inter-subject registration vs. fMRIprep’s ANTs inter-subject registration. ANTs registration has high precision for spatial normalization (Svejda et al., 2024), so one-step resampling is likely the primary explanation for OGRE’s advantages. Therefore, the one-step resampling procedure improves analytic control over the effects of individual differences in brain anatomy and head movement on fMRI analysis.

The decreased inter-individual variability with OGRE compred to fMRIprep supports the value of simultaneous registration and motion correction via one-step resampling. Inter-individual variability is a longstanding challenge in fMRI research (Dubois & Adolphs, 2016; Van Horn et al., 2008), because group averaging creates a tradeoff between statistical power and relevance to individual participants. Numerous methods exist to tackle inter-individual variability in fMRI data, but the best methods require specific data collection tools or lengthy data collection sessions (Gordon et al., 2017; Michon et al., 2022; Napadow et al., 2008), so analysis-based approaches will remain useful in many situations.

### The precision of brain extraction is not a critical driver of analysis outcomes

OGRE performs its registration and brain extraction by combining the established algorithms of FSL FLIRT and FNIRT (Andersson et al., 2007) with FreeSurfer (e.g. Fischl & Dale, 2000; Fischl et al., 2002; Fischl et al., 2004). OGRE and fMRIprep, unlike standard FreeSurfer, allow a direct comparison to FSL’s brain extraction. The OGRE pipeline led to substantially smaller brain volume estimates for task fMRI scans, but these small volumes were counter-productive: participants whose masks included more non-brain voxels had better analysis outcomes (higher Z-scores in contralateral M1), indicating that larger masks were more optimal. This is likely because spatial smoothing places information into proximal non-brain voxels, but OGRE’s HCP registration is designed for minimal preprocessing (i.e. without spatial smoothing) and thus aggressively excludes non-brain voxels – some of which contain information about a brain-surface region such as M1. Future versions of OGRE will increase brain size, which should improve OGRE’s analysis outcomes.

Otherwise, brain mask size was not correlated with analysis outcome in FSL or fMRIprep, suggesting that – despite our original motivation to remove BET brain extraction from FSL analysis – the precision of brain extraction does not necessarily play a critical role in the outcome of task fMRI analysis. Brain extraction precision may play a minor role, or a role in other metrics; but here, the inclusion of non-brain voxels did not impair analysis outcomes, likely because of the lack of task-dependent BOLD signal in distal non-brain voxels. However, even if differences in brain extraction play only a minor role in identifying task effects, Freesurfer contains numerous other advantages compared to FSL-only brain extraction, such as segmentation of gray vs. white matter.

### Use case of OGRE: improved preprocessing for researchers who prefer volumetric analysis

OGRE uses some code and concepts developed by the Human Connectome Project (HCP), including the original idea for one-step resampling; but OGRE should not be mistaken for an implementation of the HCP pipeline, which incorporates many other features including a surface-based approach to fMRI data analysis (Dickie et al., 2019; Glasser et al., 2018; Glasser et al., 2016). Surface-based analysis has many demonstrated benefits (Coalson et al., 2018), but not all analysis approaches are suitable for all research questions, groups, and protocols. Here we have quantified the advantages of Freesurfer and the one-step resampling procedure (registration and motion correction) during preprocessing, and demonstrated that those advantageous preprocessing steps can be leveraged to improve task fMRI analyses with the popular, accessible FSL toolbox.

### Limitations

Here we compared three preprocessing approaches using a motor task fMRI dataset and two traditional volumetric magnitude-based analysis methods (ROI, whole-brain). Many future directions present themselves, including using multivariate similarity-based fMRI analyses (Nili et al., 2014); and evaluating other data sets such as other tasks, task-free MRI, large datasets, and simulated data with known ground truth. Given the reduced inter-individual variability provided by OGRE preprocessing, we expect it to demonstrate benefits across a broad range of analyses and datasets, and we welcome further testing of OGRE’s robustness.

OGRE version 1.0.0 uses code derived from HCP 3.27.0, and as such may not reflect updates or bug fixes in future versions of the HCP code. Future updates to OGRE will link it to the main HCP repository to avoid this problem.

### CONCLUSION

Task fMRI analysis can be improved with preprocessing approaches that integrate tools from FSL, HCP, and FreeSurfer. We demonstrated that the “one-step resampling” preprocessing approach (i.e. simultaneous registration and motion correction, originally developed for HCP) can be applied to FSL volumetric analysis to reduce inter-individual variability throughout the brain, and increase signal in task-related regions of the brain. Gross results were largely consistent between preprocessing methods, but the one-step resampling procedure consistently led the best outcomes. Given these advantages of the one-step resampling approach, it may serve as a new methodological standard for preprocessing of fMRI data in FSL FEAT volumetric analysis. We developed the OGRE (One-step General Registration and Extraction) software to provide FSL users with an off-the-shelf general-purpose solution to implement and integrate this preprocessing approach. OGRE is available online at: https://github.com/PhilipLab/OGRE-pipeline.

## ENDING STATEMENTS

### Information sharing statement

All software is openly available at https://github.com/PhilipLab/OGRE-pipeline. The data that support the findings of this study are available on request from the corresponding author, BAP, upon reasonable request.

### Data and code availability statement

All software is openly available at https://github.com/PhilipLab/OGRE-pipeline. The data that support the findings of this study are available on request from the corresponding author, BAP, upon reasonable request.

### Author contribution statement

MPM was involved in software development (primary) and manuscript writing/editing. BAP was involved in conceptualization, software development, data analysis, funding acquisition, supervision, and manuscript writing/editing. LL and RZ were involved in data analysis and manuscript editing.

### Ethics statement

All participants gave informed consent, and all procedures were approved by the Institutional Review Board at Washington University in St. Louis School of Medicine.

